# Metabolic Collapse in Pancreatic Cancer via Combined Inhibition of Lactate Export and Thioredoxin Reductase

**DOI:** 10.1101/2025.07.09.663998

**Authors:** Adrianna Zygmunt, Eliza Turlej, Tomasz Cichoń, Marlena Golec, Magdalena Zaremba-Czogalla, Ewa Olczak, Adam Markowski, Dariusz Rakus, Jerzy Gubernator

## Abstract

Pancreatic cancer is one of the most poorly prognosed types of cancer, with a low survival rate. Cancer cells exhibit altered and rapid metabolism, and in a process known as aerobic glycolysis, they metabolize glucose to lactate even in the presence of sufficient oxygen. Altered metabolism is a hallmark of cancer, making it a promising therapeutic target. There are many potential inhibitors of cancer cell metabolism, including auranofin and syrosingopine. Auranofin induces oxidative stress in cells by generating reactive oxygen species (ROS). Meanwhile, syrosingopine is an inhibitor of the two lactate transporters, MCT1 and MCT4, which contribute to intracellular acidification. Our studies demonstrated the high biological activity of the combination of auranofin and syrosingopine against pancreatic cancer in both *in vitro* and *in vivo* models. Our proposed mechanism of action for this drug combination is based on the simultaneous induction of oxidative stress and inhibition of lactate transporters, which consequently directs cancer cells to the apoptosis pathway.

## Introduction

Pancreatic cancer, despite significant advances in technology and medicine, remains one of the deadliest types of cancer and is extremely difficult to treat. Statistics report that pancreatic cancer caused the deaths of 467,005 patients in 2022 and was the sixth leading cause of cancer-related deaths worldwide [1]. The overall five-year survival rate ranges from 2% to 9%. Furthermore, patients with inoperable tumors are expected to have a survival rate of less than five years [2]. More than 90% of all diagnosed cases of pancreatic cancer are pancreatic ductal adenocarcinoma (PDAC) [3]. This high mortality rate results from inaccurate diagnostic processes caused by the long asymptomatic progression of the disease in its early stages [4]. Potential treatment options for pancreatic cancer include surgical resection, with additional adjuvant chemotherapy improving survival rates [5]. The most applied adjuvant therapy is FOLFIRINOX, a combination of 5-fluorouracil, irinotecan, oxaliplatin, and leucovorin [6]. However, this treatment often fails to produce satisfying outcomes and is associated with numerous adverse side effects. One reason for the high resistance of pancreatic cancer to chemotherapy is the complex structure of the tumor and its microenvironment, which hinders the inward diffusion of drugs [7]. Interestingly, pancreatic cancer is the only cancer where malignant cells make up a minority of the tumor tissue (5% to 20%). The rest of the tumor consists of the surrounding microenvironment, which includes an extracellular matrix (ECM), cancer-associated fibroblasts (CAFs), pancreatic stellate cells, immune cells, endothelial cells, and soluble factors such as cytokines, chemokines, growth factors, and pro-angiogenic factors [8]. The ECM comprises various components, including collagen, fibronectin, hyaluronic acid, laminin, integrins, and matrix metalloproteinases (MMPs), which are mainly produced by cancer-associated fibroblasts and activated pancreatic stellate cells. The accumulation of ECM components disrupts the typical architecture of pancreatic tissue, leading to abnormal configurations of lymphatic and blood vessels. The stiffness of the extracellular matrix, which compresses blood vessels, may contribute to decreased blood flow, impairing effective drug delivery to cancer cells [9, 10].

Cancer cells exhibit altered metabolism, which distinguishes them from normal cells. In 1924, Otto Warburg discovered that cancer cells utilize a less efficient method of generating energy, even when sufficient oxygen is present, by breaking down glucose through a process known as aerobic glycolysis, now referred to as the Warburg effect. This metabolic change results in high lactate levels within the tumor environment [11, 12]. This may be linked to low oxygen levels and the activation of the HIF-1 transcription factor, which boosts the production of enzymes needed for glycolysis. Evidence also suggests that rapidly dividing normal cells can carry out this type of respiration [13]. Notably, about 95% of pancreatic cancers have a mutation in the KRAS gene. This gene encodes the KRAS protein, which acts as a switch, passing signals from membrane receptors to various cellular signalling pathways. Large-scale research indicates that oncogenic KRAS activation affects numerous cellular processes, promoting increased growth, movement, invasion, and survival of cancer cells by activating metabolic pathways, including MAPK, PI3K/AKT/mTOR, and NF-κB [14, 15]. Additionally, KRAS signalling enhances glucose uptake and lactate production by increasing the levels of glucose transporter-1 (GLUT-1) and several key glycolytic enzymes, including hexokinase 1 and 2 (Hk1, Hk2) and lactate dehydrogenase A (LDHA). This high rate of aerobic glycolysis provides pancreatic cancer cells with a substantial supply of glycolytic intermediates necessary for generating new cancer cells, thereby facilitating their further proliferation [16, 17].

Modified and rapid metabolism is a characteristic feature of pancreatic cancer but can also be its weakness and a therapeutic target. Nowadays, many drugs and natural substances are known to influence the metabolism of cancer cells, including syrosingopine and auranofin. Auranofin, an organic gold compound, was approved by the FDA in 1985 as an oral therapy for rheumatoid arthritis [18]. This substance has garnered interest due to its potential anticancer properties, as observed in certain tumor types, including leukemia, ovarian, lung, breast, and gastric cancers [19–23]. The probable mechanism of action of auranofin involves the inhibition of mammalian thioredoxin reductase (TrxR) in both the cytosol and mitochondria, which regulate the intracellular redox state. Auranofin-induced inhibition of thioredoxin reductase leads to an increase in ROS concentration [24]. Syrosingopine is commonly used in treating hypertension; however, recent studies suggest this drug may also possess potential anticancer properties [25]. Recent research suggests that this substance may inhibit lactate export by blocking the MCT1 and MCT4 transporters, resulting in intracellular acidification. The low pH inside the cell hampers the proper functioning of glycolytic enzymes such as phosphofructokinase 1 (PFK1), the rate-limiting enzyme. A decreased rate of aerobic glycolysis results in an insufficient energy supply to cancer cells, hindering their growth and proliferation [26, 27].

In this study, we thoroughly examined the mechanism of action of two drugs that exhibit potential impact on cancer cell metabolism – auranofin and syrosingopine, and the anticancer activity of this drug combination against pancreatic cancer. Our results indicated a strong synergistic effect between the applied drugs and high biological activity in both *in vitro* and *in vivo* studies. By understanding the mechanisms of action of both drugs, we designed a self-reinforcing therapeutic strategy that induces metabolic exhaustion in cancer cells, accompanied by severe oxidative stress, leading to mitochondrial damage and glycolysis arrest. This research proved that targeting cancer cell metabolism can be a practical approach in pancreatic cancer treatment.

## Results

### Cytotoxicity of auranofin and syrosingopine combination towards pancreatic cancer cells

To assess the anticancer potential of auranofin and syrosingopine combination, we examined the cytotoxic activity against two human pancreatic adenocarcinoma cell lines (AsPC-1 and BxPC-3) and a mouse pancreatic cancer cell line (KPCY) *in vitro*. As a control cell line, we used normal human dermal fibroblasts (NHDF). Cells were treated with free auranofin, syrosingopine, or a combination of both compounds in a 1:1 molar ratio at concentrations ranging from 0.5 to 10 µM for 24 and 48 hours. DMSO was used as a solvent control in a volume corresponding to the drug concentration. Cell viability was established using the colorimetric SRB test, which is based on the ability of the SRB dye to electrostatically bind basic amino acids in cells. The number of bound proteins directly determines the number of viable cells in culture [28]. All the cancer cell lines were sensitive to the combination therapy, depending on the dose used and incubation time (see Supplementary Materials, **Figures S1 and S2**). More importantly, syrosingopine significantly enhanced the anticancer effect of auranofin. Furthermore, the drug combination appeared to be less toxic to the reference cells. Based on the cell survival curves, we determined the IC50 values, which are summarized in **Table 1**. To confirm the synergistic effect between the tested compounds, we also determined the combination index (CI). The CI values are presented in **Table 2**.

**Table 1.**
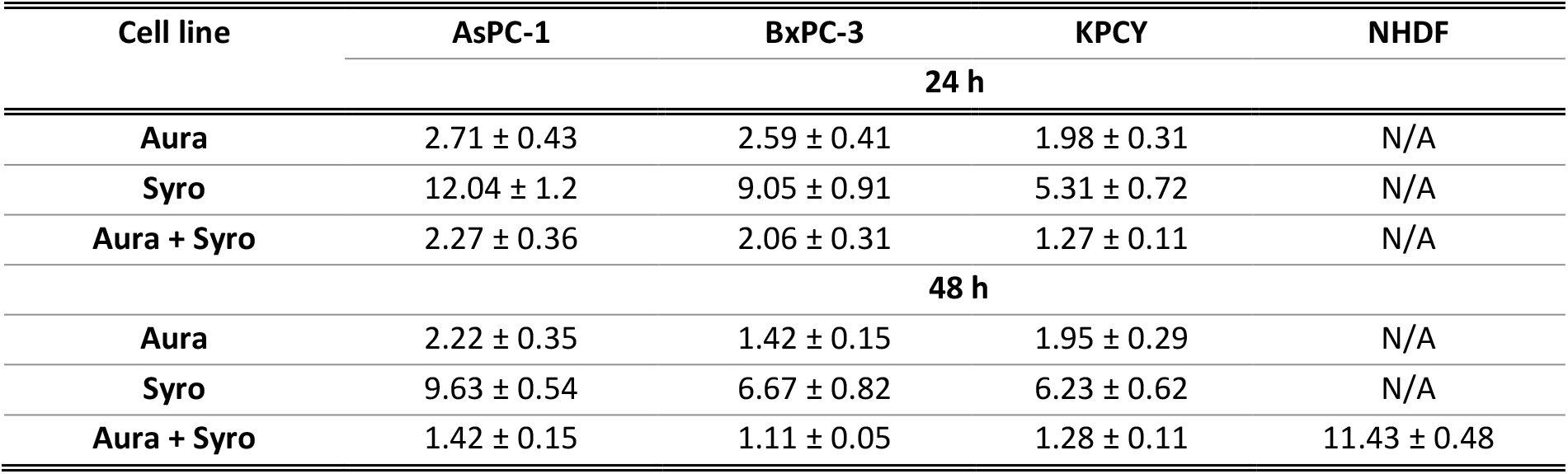
IC50 values (µM) of free compounds and drug combinations for the tested cell lines. Explanations of abbreviations used: Aura – auranofin; Syro – syrosingopine

| Cell line | AsPC-1 | BxPC-3 | KPCY | NHDF |
| --- | --- | --- | --- | --- |
| 24 h |  |  |  |  |
| Aura | 2.71 ± 0.43 | 2.59 ± 0.41 | 1.98 ± 0.31 | N/A |
| Syro | 12.04 ± 1.2 | 9.05 ± 0.91 | 5.31 ± 0.72 | N/A |
| Aura + Syro | 2.27 ± 0.36 | 2.06 ± 0.31 | 1.27 ± 0.11 | N/A |
| 48 h |  |  |  |  |
| Aura | 2.22 ± 0.35 | 1.42 ± 0.15 | 1.95 ± 0.29 | N/A |
| Syro | 9.63 ± 0.54 | 6.67 ± 0.82 | 6.23 ± 0.62 | N/A |
| Aura + Syro | 1.42 ± 0.15 | 1.11 ± 0.05 | 1.28 ± 0.11 | 11.43 ± 0.48 |

**Table 2.**
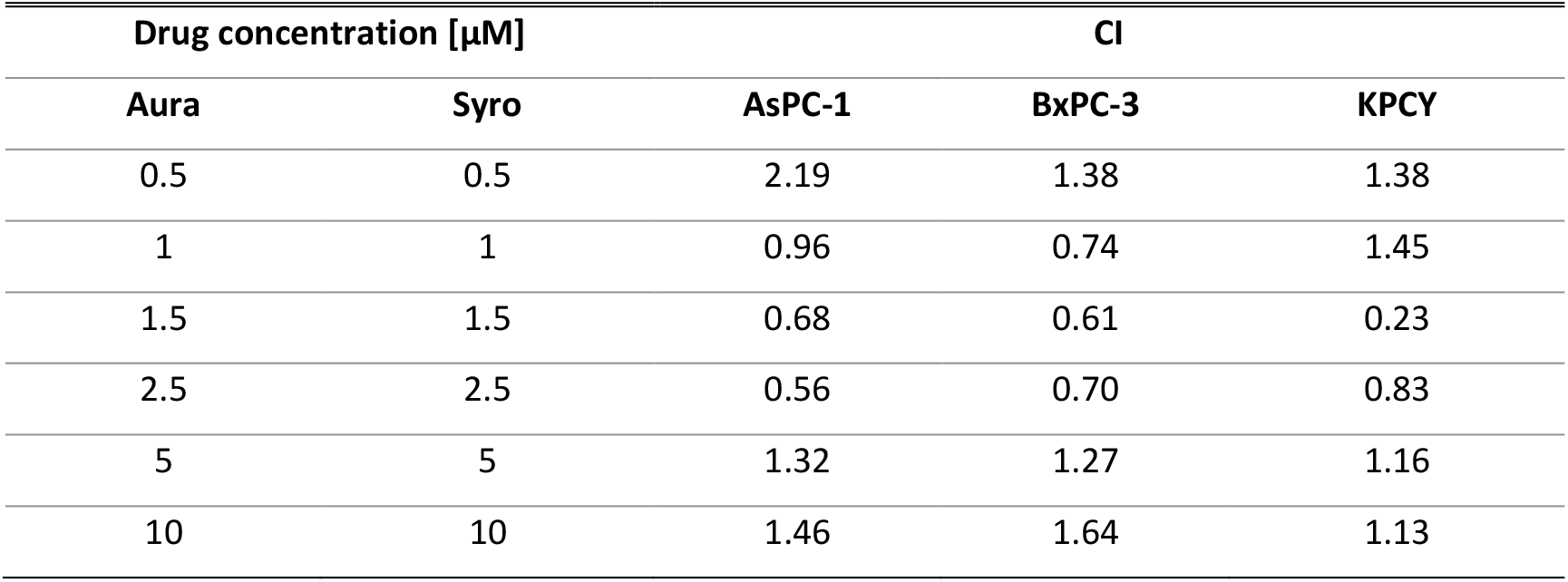
Determined CI values based on the SRB test performed after 48-hour incubation of cells with test compounds. Interpretation of results: CI < 1 synergism; CI = 1 additive; CI > 1 antagonism. Explanations of abbreviations used: Aura – auranofin; Syro – syrosingopine

| Drug concentration [μM] | | CI | | |
| --- | --- | --- | --- | --- |
| Aura | Syro | AsPC-1 | BxPC-3 | KPCY |
| 0.5 | 0.5 | 2.19 | 1.38 | 1.38 |
| 1 | 1 | 0.96 | 0.74 | 1.45 |
| 1.5 | 1.5 | 0.68 | 0.61 | 0.23 |
| 2.5 | 2.5 | 0.56 | 0.70 | 0.83 |
| 5 | 5 | 1.32 | 1.27 | 1.16 |
| 10 | 10 | 1.46 | 1.64 | 1.13 |

To confirm the data obtained in the SRB assay, we determined the level of secreted lactate dehydrogenase (LDH), an enzyme that serves as a marker for cytotoxicity tests. The LDH level was elevated in human pancreatic cancer cells using the LDH-Glo™ Cytotoxicity Assay, as per the manufacturer’s instructions. AsPC-1 and BxPC-3 cells were incubated for 48 hours with 1.5 and 2.5 µM auranofin, syrosingopine, and the drug combination. The outcomes posted on **Figure 1** show a significant increase in LDH levels in the culture medium for both pancreatic cancer lines after treatment with auranofin and dual therapy. Thus, the results obtained confirm the high biological activity of the combination of auranofin and syrosingopine against pancreatic adenocarcinoma cells.

**Figure 1.**
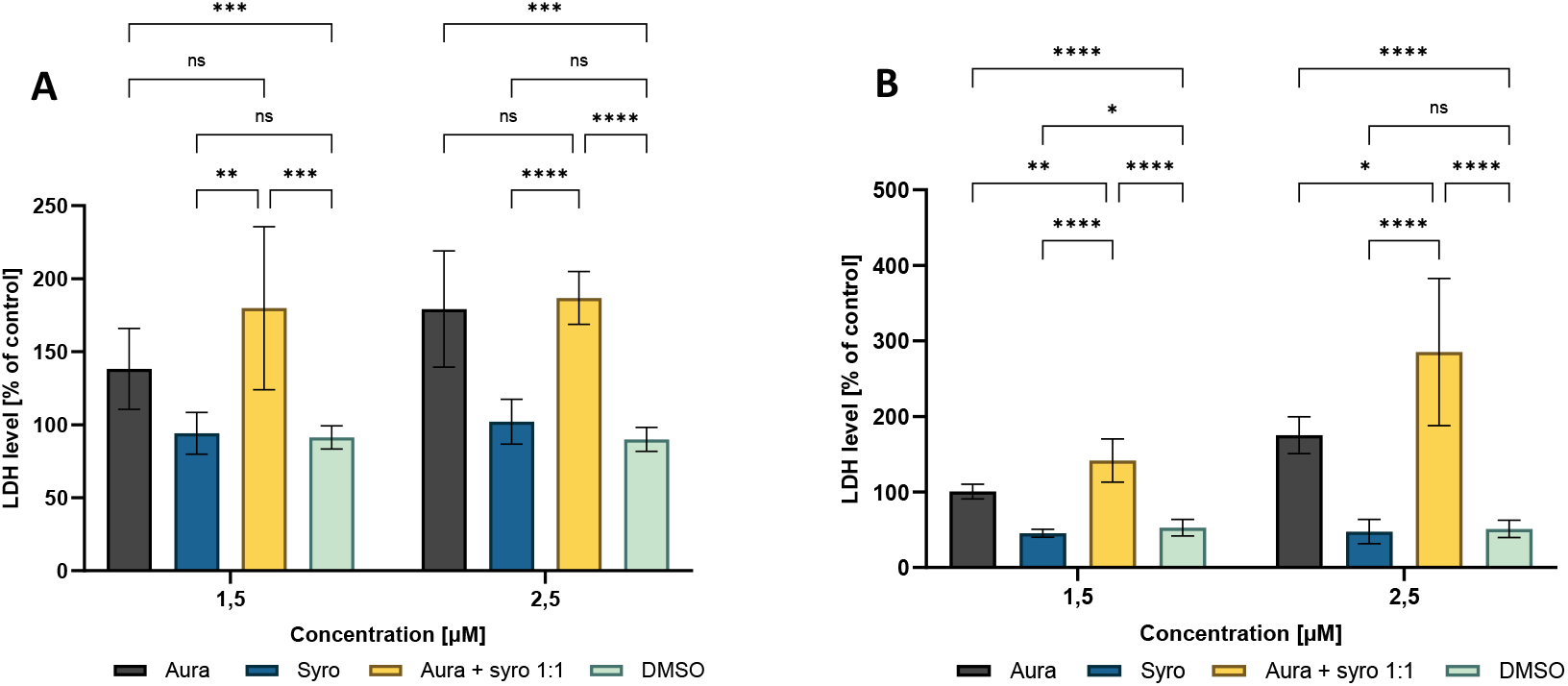
Level of released LDH from AsPC-1 (A) and BxPC-3 (B) cells after 48-hour incubation with auranofin, syrosingopine, drug combination at 1:1 molar ratio, and DMSO. Untreated cells were a 100% normalized control. LDH levels after treatment with test compounds were expressed as % of the control. Bar graphs show mean values with standard deviation from three independent biological replicates. Statistical significance was calculated using a two-way ANOVA test (Tukey’s modification). ^****^ P < 0.0001; ^***^ P < 0.0002; ^**^ P < 0.0021; ^*^P < 0.033; ns - not statistically significant (0.1234). Explanations of abbreviations used: Aura - auranofin, Syro - syrosingopine

### A combination of auranofin and syrosingopine inhibits aerobic glycolysis in human pancreatic cancer cells

To investigate whether the combination of auranofin and syrosingopine inhibits aerobic glycolysis in human pancreatic cancer cells, we measured lactate levels in the medium, determined cellular ATP levels, and assessed hexokinase 2 (HK-2) expression. According to available literature, syrosingopine is an inhibitor of lactate transporters (MCTs), which are responsible for removing excess lactate from cells, among other functions. Dysfunction of these transporters can lead to excessive lactate accumulation within cells and severe intracellular acidification, which in turn disrupts metabolic pathways. We determined the lactate level in the medium collected from the cells after a 48-hour incubation with 1.5 and 2.5 µM auranofin, syrosingopine, and the drug combination using the Lactate-Glo Assay according to the manufacturer’s instructions. We observed a decrease in lactate levels in both the primary pancreatic cancer line and the metastatic line after treating the cells with a dual therapy (**Figures 2A and 2B**). These results suggest inhibition of MCT transporters’ activity and lactate accumulation in cancer cells. Since excessive lactate accumulation leads to the acidification of the intracellular environment and the inhibition of several glycolytic enzymes, we correlated the results obtained with the level of intracellular ATP, which is the final product of the glycolysis process. To evaluate the ATP level in pancreatic cancer cells, we performed an analogous experiment to the lactate assay using the CellTiter-Glo 2.0 Cell Viability Assay. As expected, the level of endogenous ATP was comparably reduced for both cancer cell lines after treatment with the tested drug combination in a dose-dependent manner (**Figure 2C and 2D**). The results obtained not only confirm the cytotoxicity of the proposed drug combination but also indicate disruption of the glycolysis process in cancer cells. To support our hypothesis regarding the inhibition of glycolysis by the proposed drug combination, we additionally examine the expression of hexokinase 2 in pancreatic cancer cells. HK2 is an isoform of an enzyme prevalent in cancer cells that is responsible for catalysing the first step of aerobic glycolysis. Hence, HK2 is considered one of the most essential enzymes regulating the rate of glycolysis. Inhibition of this enzyme’s activity can prevent cells from continuously extracting energy, which is critical for rapid cell proliferation. The expression level of hexokinase 2 was determined in pancreatic cancer cells after 48 hours of treatment with the combination of auranofin and syrosingopine. Analysis of the results revealed reduced expression of HK-2, primarily in BxPC-3 cells (**Figure 2F**). Interestingly, we did not observe a comparable effect in the AsPC-1 cell line (**Figure 2E**). Nevertheless, our study proved that one of the mechanisms of action of the proposed drug combination is likely to be interference with the aerobic glycolysis pathway in cancer cells.

**Figure 2.**
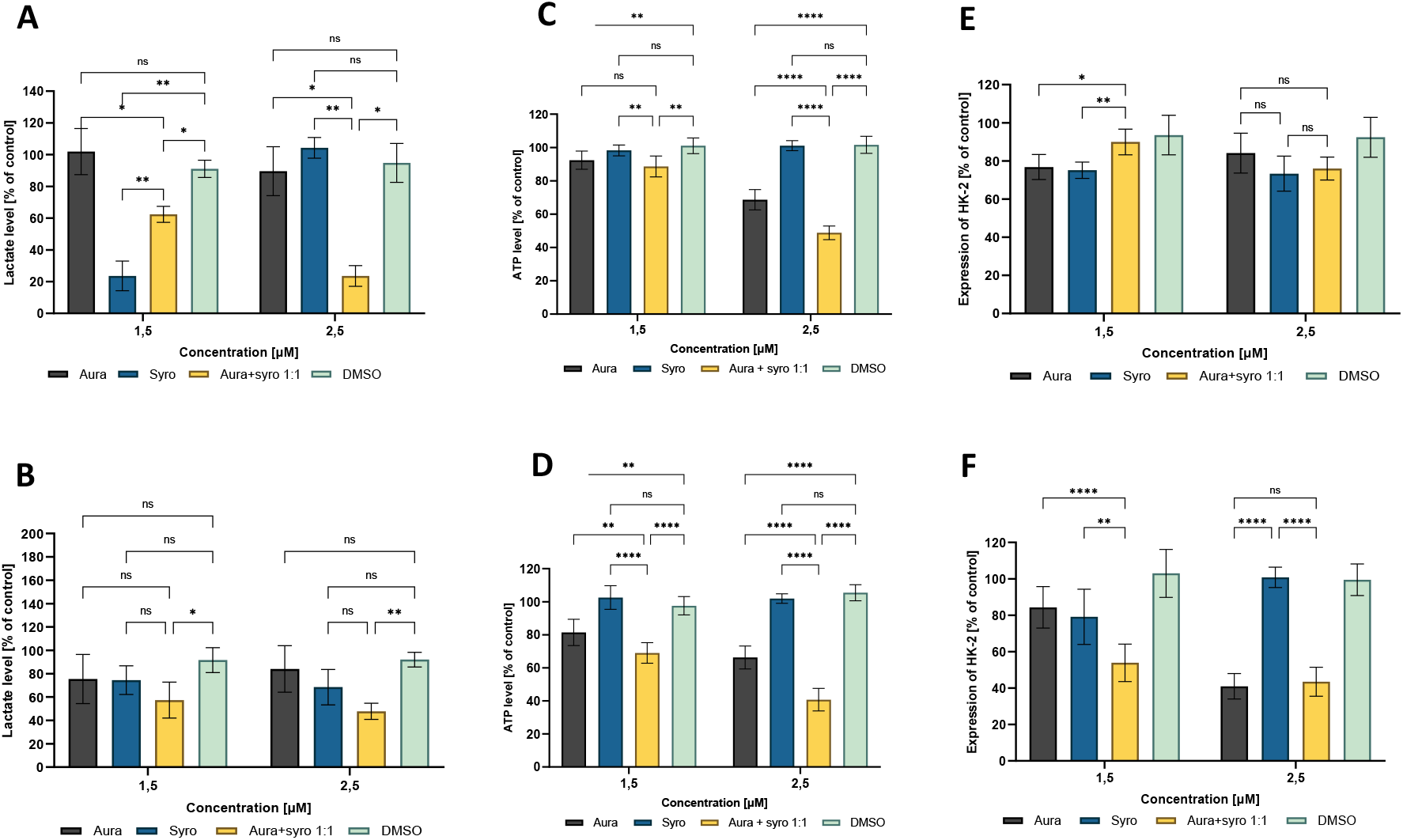
Lactate level (A-B), intracellular ATP level (C-D), and HK-2 expression (E-F) in AsPC-1 (upper) and BxPC-3 (bottom) cells after 48-hour incubation with auranofin, syrosingopine, drug combination at 1:1 molar ratio, and DMSO. Untreated cells were a 100% normalized control. LDH levels after treatment with test compounds were expressed as % of the control. Bar graphs show mean values with standard deviation from three independent biological replicates. Statistical significance was calculated using a two-way ANOVA test (Tukey’s modification). ^****^ P < 0.0001; ^***^ P < 0.0002; ^**^ P < 0.0021; ^*^P < 0.033; ns - not statistically significant (0.1234). Explanations of abbreviations used: Aura - auranofin, Syro – syrosingopine

### A combination of auranofin and syrosingopine generates oxidative stress in human pancreatic cancer cells

The mechanism of action of auranofin is based, among other factors, on the generation of free radicals through the inhibition of thioredoxin reductase, an essential enzyme involved in regulating the cellular redox state. By blocking the activity of this enzyme, oxidative stress within the cell increases, which can potentially lead to cell death. Therefore, the following research focused on assessing the level of oxidative stress in pancreatic cancer cells following treatment with a drug combination. To gain a broader perspective, we first measured ROS levels in the cancer cells and then correlated these results with glutathione (GSH) levels. GSH is regarded as one of the most crucial endogenous antioxidant systems for cells, playing a key role in maintaining redox homeostasis and protecting cells from oxidative damage. Changes in glutathione levels can be triggered by increased free radicals, indicating elevated oxidative stress within the cells. We measured ROS and GSH levels in pancreatic cancer cells after 48 hours of incubation with a combination of auranofin and syrosingopine using the ROS-Glo™ H2O2 Assay and the GSH-Glo™ Glutathione Assay, respectively. The results, presented in **Figure 3**, provided several insights. Firstly, we demonstrated that auranofin’s mechanism of action in pancreatic cancer cells also involves the generation of free radicals. Secondly, syrosingopine significantly enhances the effects of auranofin, even though it does not generate ROS in cancer cells itself. Finally, we found that increased levels of ROS correspond to reduced levels of GSH in the cells, indicating a distinct oxidative stress prevailing within the pancreatic cancer cells.

**Figure 3.**
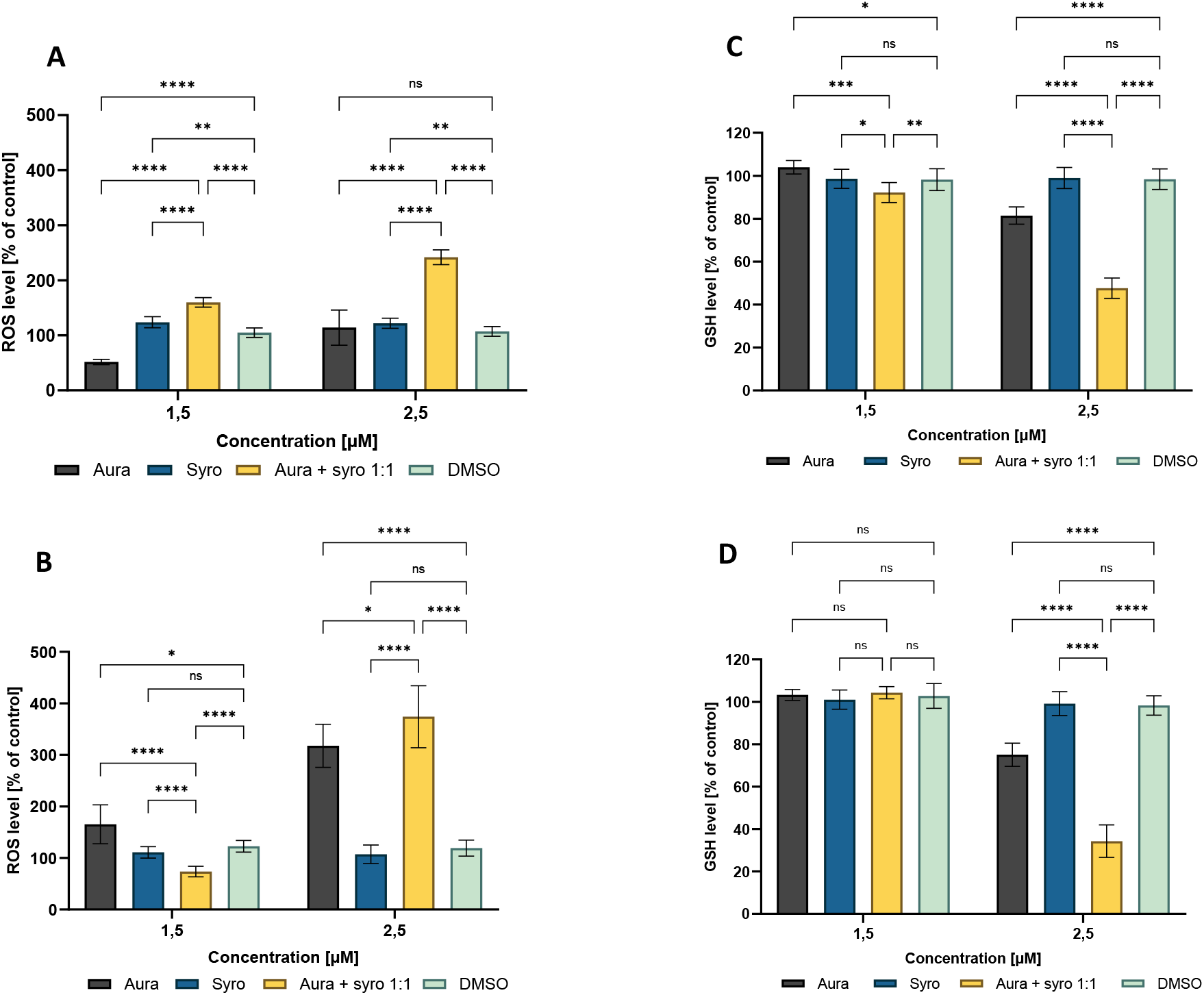
ROS (A-B) and GSH (C-D) levels in AsPC-1 (upper) and BxPC-3 (bottom) cells after 48-hour incubation with auranofin, syrosingopine, drug combination at 1:1 molar ratio, and DMSO. Untreated cells were a 100% normalized control. LDH levels after treatment with test compounds were expressed as % of the control. Bar graphs show mean values with standard deviation from three independent biological replicates. Statistical significance was calculated using a two-way ANOVA test (Tukey’s modification). ^****^ P < 0.0001; ^***^ P < 0.0002; ^**^P < 0.0021; ^*^ P < 0.033; ns - not statistically significant (0.1234). Explanations of abbreviations used: Aura - auranofin, Syro – syrosingopine

### Auranofin and syrosingopine combination induces apoptosis in human AsPC-1 and BxPC-3 cell lines

In the following step, we investigated whether a combination of auranofin and syrosingopine induces apoptosis in pancreatic cancer cells. We estimated the progress of apoptosis with the RealTime-Glo™ Annexin V Apoptosis and Necrosis Assay reagent kit. This kinetic assay measures the exposure of phosphatidylserine on the cell membrane surface in real time. Annexin V binding in cells is detected using a luminescent signal, while necrosis detection is based on a fluorescent signal induced by a DNA dye. We monitored the progression of apoptosis and necrosis in AsPC-1 and BxPC-3 cells for 48 hours of drug treatment, and the results are presented in **Figure 4**. In both the primary pancreatic cancer line and the metastatic line, a significant signal from annexin V is observed after treatment with the combination of auranofin and syrosingopine. We perceived the induction of apoptosis as early as 6 hours in the case of BxPC-3 cells, while in the case of the AsPC-1 line, it occurred after 12 hours. Additionally, we assessed mitochondrial membrane potential to determine if activation of the apoptosis pathway is associated with mitochondrial damage. Changes in mitochondrial membrane potential we measured by JC-1 staining after 48 hours of treatment with the tested compounds. Changes in membrane potential characterized both pancreatic cancer cell lines; however, auranofin alone decreased mitochondrial potential to a greater extent than the drug combination (**Figure 4**). These results indicate that auranofin, in particular, causes mitochondrial damage.

**Figure. 4.**
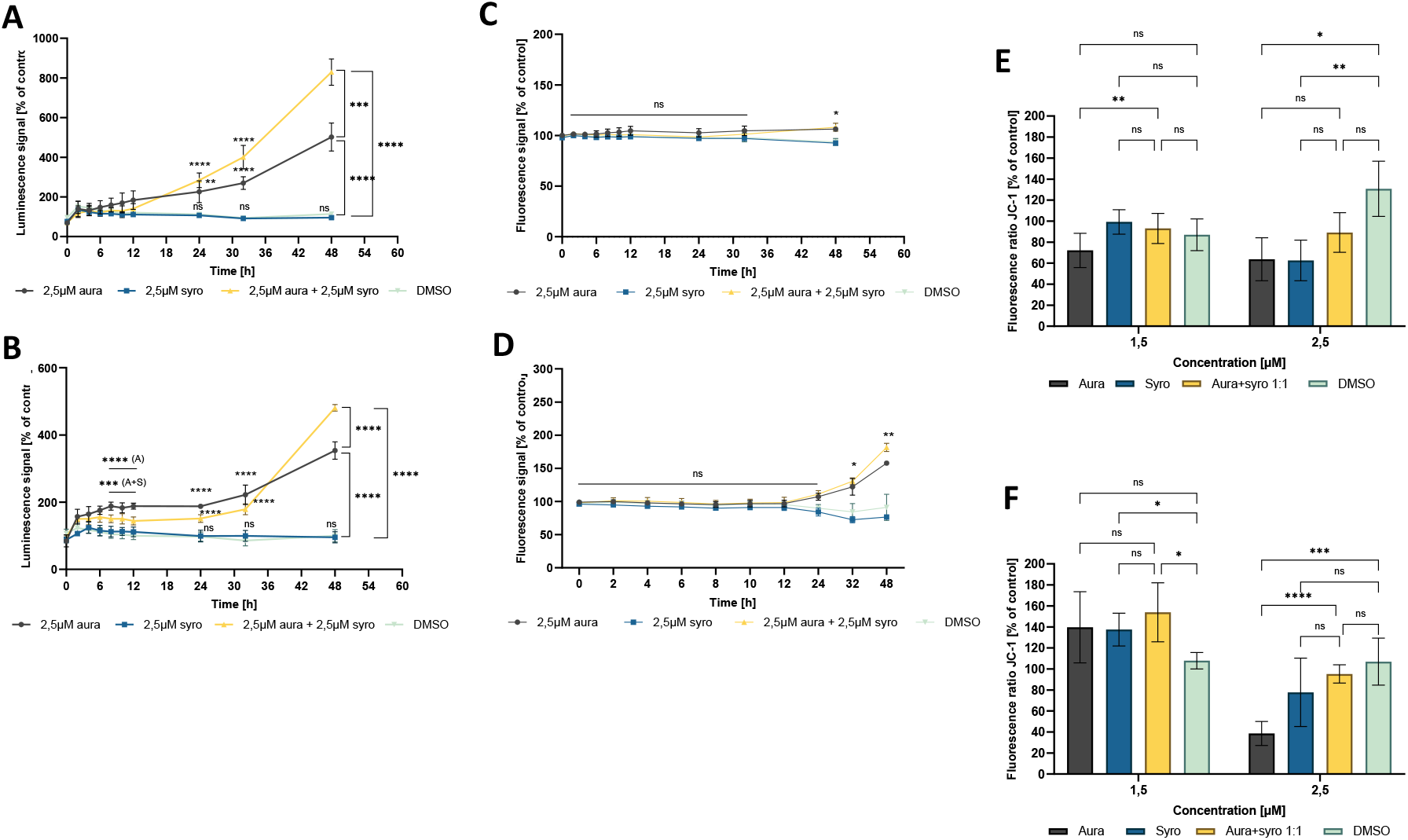
Real-time apoptosis (A-B), necrosis (C-D), and mitochondrial membrane potential (E-F) in AsPC-1 (upper) and BxPC-3 (bottom) cells after 48-hour incubation with auranofin, syrosingopine, drug combination at 1:1 molar ratio, and DMSO. Untreated cells were a 100% normalized control. LDH levels after treatment with test compounds were expressed as % of the control. Bar graphs show mean values with standard deviation from three independent biological replicates. Statistical significance was calculated using a two-way ANOVA test (Tukey’s modification). ****P < 0.0001; ***P < 0.0002; **P < 0.0021; *P < 0.033; ns - not statistically significant (0.1234). Explanations of abbreviations used: Aura - auranofin, Syro – syrosingopine

### Drug combination activity against a 3D model of pancreatic cancer

We also evaluated the biological activity of the proposed drug combination against a 3D culture (spheroid) of BxPC-3 cells. Spheroids are 3D cell cultures that provide a more accurate model reflecting tumor conditions than traditional 2D cultures. Hence, studies using spheroids can provide more precise information about the cytotoxicity of test compounds. As a 3D model line, we selected BxPC-3 cells and treated the cultured triplicate cultures with a combination of drugs at appropriate concentrations for 72 hours. We used microscopic imaging to assess morphological changes in the spheroids, and viability was determined by bioluminescent measurements of ATP levels in cultures. Microscopic imaging (**Figure 5**) showed disturbed morphology of the spheroids after incubation with the combination of auranofin and syrosingopine. In addition to an apparent envelope presumably composed of dead cells, changes were observed in the density of the spheroid core compared to the control. Next, we evaluated the cytotoxicity of the drug combination against 3D cultures using the CellTiter-Glo 3D Assay. ATP levels were significantly reduced after treating spheroids with the combination of compounds, which at a concentration of 2.5 μM decreased viability levels by about 50%. Free auranofin had a weaker effect against the 3D model, while syrosingopine showed no impact on spheroid viability (**Figure 5**).

**Figure 5.**
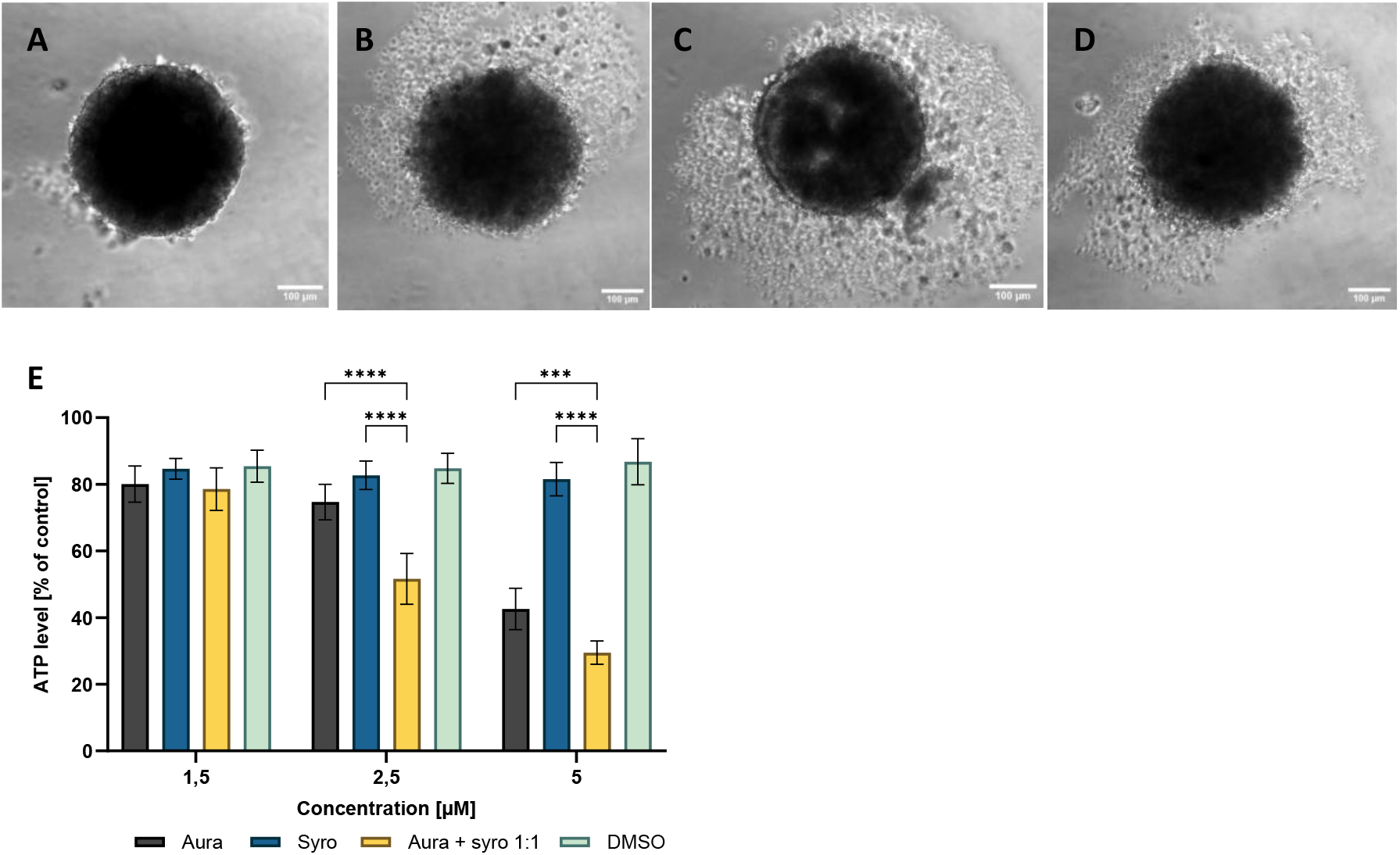
Effect of auranofin and syrosingopin combination on morphology and viability of spheroids after 72 h of treatment. A - untreated spheroids (control), B - 5 μM auranofin, C - 5 μM syrosingopin, D - aura + syro combination 1:1. Representative microscopic images were taken with a Zeiss Colibri microscope. E - ATP levels in 3D cultures of BxPC-3 lines after 72-hour incubation with auranofin, syrosingopine, drug combination at 1:1 molar ratio, and DMSO. Untreated cells were a 100% normalized control. LDH levels after treatment with test compounds were expressed as % of the control. Bar graphs show mean values with standard deviation from three independent biological replicates. Statistical significance was calculated using a two-way ANOVA test (Tukey’s modification). ^****^ P < 0.0001; ^***^ P < 0.0002; ^**^ P < 0.0021; ^*^ P < 0.033; ns - not statistically significant (0.1234). Explanations of abbreviations used: Aura - auranofin, Syro – syrosingopine

### Dual therapy impedes pancreatic tumor growth in an *in vivo* study

To evaluate the anticancer potential of auranofin and syrosingopine combination, we performed an *in vivo* study using a mouse model of pancreatic cancer, C57BL/6NCrL. The drugs were injected into the tumors of mice inoculated with KPCY mouse pancreatic cancer cells. The mice were divided into five experimental groups. The drug dose used was 5mg/kg body weight and was administered alone and in combination from days 14 to 18 of the experiment. As a solvent control, we used 30% DMSO + 30% PEG400. We measured both the tumor volume and body weight of the mice at specific time intervals. The results presented in **Figure 6** did not show inhibition of KPCY tumor growth when auranofin and syrosingopine were administered separately. In contrast, administration of the dual therapy to mice at a dose of 5 mg/kg body weight resulted in a reduction in tumor volume of almost 50% by day 16 of the experiment. Notably, there was no increase in tumor growth kinetics after discontinuation of the treatment. Based on the data obtained, we concluded that the combination of auranofin and syrosingopin significantly inhibits the growth and development of pancreatic cancer tumors in an animal model. We also observed no change in the body weight of the mice during the experiment, suggesting no toxic effect of dual therapy.

**Figure 6.**
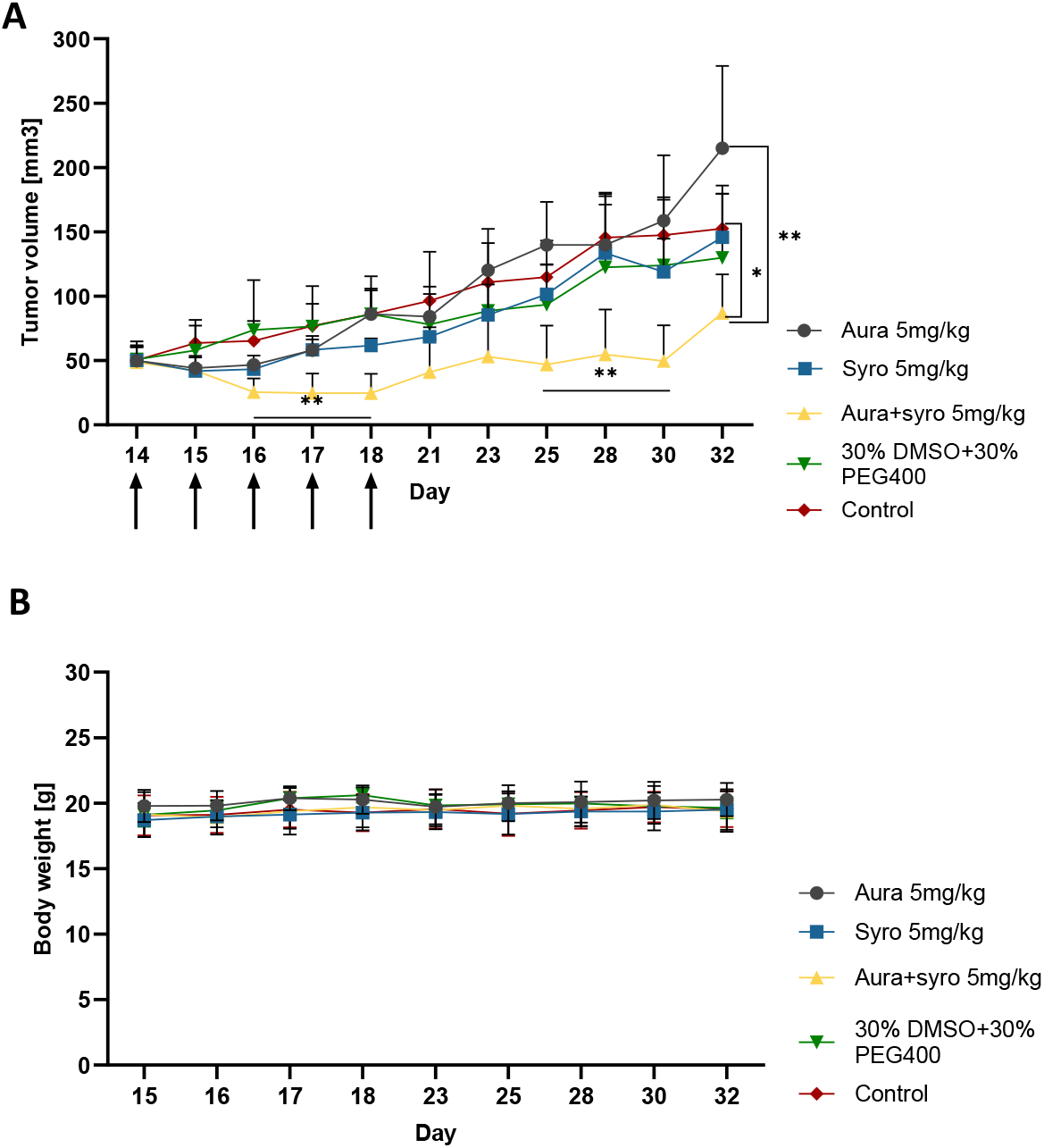
Kinetics of KPCY tumor growth (A) and body weight changes in mice (B) throughout the experiment after administration of auranofin (5 mg/kg), syrosingopin (5 mg/kg), drug combination (5 mg/kg aura + 5 mg/kg syro) and solvent control (30% DMSO + 30% PEG400). The control group of the experiment was untreated mice. Line graphs display mean values with standard deviations from five biological replicates. Arrows indicate days of administration of the formulations. Explanations of abbreviations used: Aura - auranofin, Syro – syrosingopine

## Discussion

Despite significant advances in medicine over the past few decades, pancreatic cancer remains one of the most challenging malignancies to treat. It continues to be one of the deadliest cancer types worldwide. The low survival rate among patients is primarily due to difficulties in early diagnosis and the highly invasive nature of the disease [29]. In its early stages, pancreatic cancer is often asymptomatic, and diagnosis typically occurs only after the tumor has metastasized to other organs. As a result, late detection severely limits treatment options, which are usually reduced to palliative care. Current therapeutic approaches offer minimal hope for a complete cure, particularly for patients in advanced stages of the disease. Treatment generally involves partial resection of the pancreas, which depends largely on the tumor’s location, and chemotherapy that extends survival by only a few months. Notably, only a small proportion of patients are eligible for advanced therapies. Another major challenge is the high resistance of pancreatic tumors to available drugs [30]. Therefore, there is a pressing need to search for new anticancer compounds continually and to deepen our understanding of pancreatic cancer biology. As previously mentioned, cancer cells exhibit altered metabolism and require substantial amounts of energy to sustain rapid proliferation. A well-known phenomenon observed in many tumors, known as the Warburg effect, involves a metabolic shift toward aerobic glycolysis rather than oxidative phosphorylation. This adaptation enables cancer cells to rapidly convert glucose into energy in the form of ATP, while also producing essential building blocks for cell growth, such as nucleotides and lipids [31, 32]. Therefore, targeting the unique metabolic features of cancer cells to inhibit their growth and survival seems to be a promising approach to treating pancreatic cancer. In this research, we focused on two potential inhibitors of cancer cell metabolism: auranofin and syrosingopine. In recent years, the anticancer activity of auranofin has been demonstrated in several studies across various cancer types, including breast, gastric, ovarian, and lung cancers, as well as leukemia [20–24]. Emerging research also highlights its potential use in treating pancreatic cancer, indicating growing interest in this compound as a prospective anticancer agent [33]. On the other hand, syrosingopine remains relatively understudied, and its anticancer potential is only beginning to be explored, with preliminary investigations conducted in models of leukemia, lung cancer, and breast cancer [34,35]. Our main goal was to discover the mechanism of action of the combination of these two compounds in the context of pancreatic adenocarcinoma. We first examined the biological activity of this drug combination against two human pancreatic adenocarcinoma lines: primary BxPC-3 and metastatic AsPC-1, as well as the mouse line KPCY. We chose human dermal fibroblasts (NHDF) as the reference line. The SRB assay demonstrated that the combination of auranofin and syrosingopine significantly enhanced cancer cell death compared to either compound alone in all tested cancer cell lines. In contrast, normal cells exhibited a much lower mortality rate. These findings were further supported by LDH release measurements, indicating reduced cell viability after treatment in AsPC-1 and BxPC-3 cells. Moreover, the combination index (CI), calculated using the Chou-Talalay method, confirmed a strong synergistic effect between auranofin and syrosingopine. Combination therapies are increasingly being adopted in cancer treatment. Beyond the clear benefit of enhanced therapeutic efficacy, this approach offers the important advantage of dose reduction, which can potentially minimize toxicity and side effects. Several studies have reported synergistic effects of auranofin when combined with other compounds. For instance, in breast cancer models, the co-administration of auranofin and meclofenoxate (mezupron) resulted in stronger inhibition of cancer cell growth *in vitro* [22]. A similar synergistic effect was observed with auranofin and ascorbic acid, confirmed both *in vitro* and in a mouse model [36]. Han et al. also demonstrated synergy between auranofin and celecoxib in colorectal cancer cells. However, in these studies, although auranofin was used at relatively low concentrations (≤1 μM), significant anticancer effects required much higher doses of the second compound, at least 10 μM for mezupron and celecoxib, and 1 mM for ascorbic acid [37]. A similar pattern was observed in leukemia and multiple myeloma models treated with a combination of syrosingopine and metformin, where effective concentrations were 5 μM and 5 mM, respectively [26]. In contrast, the results presented in this study are promising: a concentration of just 1.5 μM for both auranofin and syrosingopine was sufficient to strongly inhibit the proliferation of pancreatic cancer cells. This suggests that syrosingopine may enhance the anticancer effects of auranofin more effectively than the previously studied agents. The mechanism of action of the auranofin and syrosingopine combination was investigated in two human pancreatic cancer cell lines: AsPC-1 and BxPC-3. Based on the higher cytotoxic effect observed after 48 hours, the same incubation time was used for mechanistic studies. The first experiment focused on measuring lactate levels in the culture medium. Lactate, a key product of aerobic glycolysis, supports tumor growth and is actively exported by MCT transporters to prevent intracellular acidification. Blocking MCT activity can lead to lactate accumulation, reduced glycolytic flux, and decreased ATP and NADPH levels [38, 39]. Previous studies by Benjamin et al. showed that syrosingopine reduces extracellular lactate levels, including in HeLa cervical cancer cells [26]. In our research, a noticeable reduction in lactate levels was observed following treatment with the drug combination. These results were further supported by measurements of intracellular ATP and HK-2 expression in the cells. Analysis of the results revealed a low level of ATP and a relatively reduced level of HK-2 expression in AsPC-1 and BxPC-3 cells, particularly after treatment with the combination of auranofin and syrosingopine, which was dependent on the concentration of the drugs. The data obtained may indicate impaired aerobic glycolysis in cancer cells. Redox balance plays a vital role in the survival and progression of cancer cells. Excessive production of reactive oxygen species (ROS) can damage essential cellular components, including proteins, lipids, and DNA [40]. Given the vulnerability of cancer cells to oxidative stress, therapeutic strategies that induce elevated ROS levels hold significant potential. Studies conducted on gastric and lung cancer cell models have demonstrated that auranofin increases intracellular ROS, leading to reduced cell viability. A similar effect was anticipated in pancreatic cancer cells. Our research confirmed the effect of auranofin, showing a marked increase in ROS levels in both human pancreatic cancer cell lines. Interestingly, the combination of auranofin and syrosingopine proved to be even more effective, despite syrosingopine alone having no significant impact on ROS levels. Elevated ROS levels are typically associated with a reduction in intracellular glutathione. As expected, the GSH level was decreased in AsPC-1 and BxPC-3 cells, indicating an increase in oxidative stress. Our next study was designed to test whether elevated levels of ROS induce apoptosis in pancreatic cancer cells. This process is regulated by two main pathways: the intrinsic (mitochondrial) and the extrinsic (receptor-mediated) apoptotic pathways. Apoptosis is most triggered via the intrinsic pathway, typically in response to DNA damage or oxidative stress, which activates members of the Bcl-2 protein family. When pro-apoptotic proteins (such as Bax and Bak) outweigh antiapoptotic ones (like Bcl-2 and Bcl-XL), mitochondrial membrane permeability increases, leading to the release of cytochrome c into the cytoplasm. There, cytochrome c binds to Apaf-1, forming a complex that activates caspase-9, followed by caspases-3 and -7, ultimately resulting in programmed cell death [41, 42]. Our studies conducted on pancreatic cancer cell lines confirmed that the tested drug combination was capable of inducing apoptosis. Auranofin alone also contributed to the progression of this process. These results were further supported by assessment of mitochondrial membrane potential. Mitochondrial membrane potential is a key indicator of mitochondrial health, and its reduction often signifies membrane damage, which can lead to the release of cytochrome c and the initiation of apoptosis. One possible cause of such changes is an increase in intracellular ROS levels. The results obtained in this study supported the initial hypothesis, revealing dose-dependent alterations in membrane potential. The proposed mechanism of action for the combination of auranofin and syrosingopine in pancreatic cancer cell lines involves the induction of oxidative stress. Auranofin contributes to this by generating reactive oxygen species (ROS) and inhibiting enzymes responsible for detoxifying free radicals. In response to redox imbalance, cancer cells activate defence mechanisms, including a sharp acceleration of glycolysis to produce ATP and NADPH. However, the excess ROS damages mitochondria, impairing oxidative phosphorylation as a source of ATP. To compensate, the cells increase glycolytic activity, resulting in elevated lactate production. Normally exported from the cell, lactate accumulates due to syrosingopine-mediated inhibition of monocarboxylate transporters. This leads to intracellular acidification, which disrupts the function of glycolytic enzymes. As a result, glycolysis is suppressed, triggering apoptosis and ultimately leading to cancer cell death [43, 44]. We also evaluated the anticancer potential of the proposed drug combination using a more complex research model of pancreatic cancer, which included 3D cultures and *in vivo* studies. The combination of auranofin and syrosingopine disrupted the spheroid structure, and in addition, the measured ATP levels indicated low viability of the cultured cells. Moreover, *in vivo* studies demonstrated that the combination of auranofin and syrosingopine may be more effective than either compound alone. Specifically, tumor growth was significantly inhibited following combination treatment, and this suppression remained more stable after treatment cessation compared to tumors treated with auranofin or syrosingopine alone. Additionally, monitoring the animals’ body weight throughout the experiment suggests that the dosage of 5 mg/kg body weight is well-tolerated, with no notable deviations or signs of compound-related toxicity. Based on these findings, the combined use of auranofin and syrosingopine appears promising for future therapeutic strategies against pancreatic cancer.

## Materials and Methods

### Materials

Auranofin (≥ 95% purity) was purchased from Cayman Chemicals (Michigan, USA). Syrosingopine was purchased from Biosynth (Bratislava, Slovakia). Sulforhodamine B (SRB) and TRIS (hydroxymethyl)aminomethane were obtained from Sigma Aldrich (Life Science®, Saint Louis, MO, USA). Trichloroacetic acid (TCA), acetic acid, and dimethylsulfoxide (DMSO) were sourced from Chempur (Piekary Slaskie, Poland). PBS pH 7.4 was acquired from EURx Molecular Biology Products (Gdansk, Poland). RPMI 1640 and MEMα cell culture media were obtained from BioWest (Nuaillé, France). Fetal bovine serum (FBS), 100x antibiotic-antimycotic, and glutamine were sourced from BioWest (Nuaillé, France). CellTiter-Glo® 2.0 Cell Viability Assay, CellTiter-Glo® 3D Cell Viability Assay, GSH-Glo_™_ Glutathione Assay, ROS-Glo_™_ H2O2 Assay, LDH-Glo_™_ Cytotoxicity Assay, RealTime-Glo_™_ Annexin V Apoptosis and Necrosis Assay, Lactate-Glo_™_ Assay, and Glucose-Glo_™_ Assay were purchased from Promega GmbH (Walldorf, Germany). Agarose standard was acquired from Carl Roth GmbH (Karlsruhe, Germany).

### Cell Culture

Human pancreatic cell lines AsPC-1 (derived from metastatic pancreatic ductal adenocarcinoma, PDAC) and BxPC-3 (derived from a primary tumor) were purchased from ATCC (Manassas, VA, USA). The cells were cultured in RPMI-1640 medium supplemented with 10% fetal bovine serum, 1% glutamine, and 1% antibiotic-antimycotic. The mouse pancreatic cell line KPCY (Kerafast, Inc., Boston, MA, USA) was maintained under the same conditions as the human pancreatic cell lines. The reference cells, NHDF (normal human dermal fibroblasts), were cultured in MEMα medium supplemented with the same additives. All the cell lines were maintained at 37° C in an incubator with 5% CO2 (Memmert ICO150 Med., Memmert, Buchenbach, Germany).

### Cytotoxicity Assay

The biological activity of the tested drugs was evaluated using the SRB cytotoxicity assay (ref.). Cells were seeded into 96-well plates with the appropriate culture medium and incubated for 24 hours. The number of KPCY cells was 3 × 10^3^ per well, while the other cell lines were seeded at 4.5 × 10^3^ per well. The cells were treated with different concentrations of auranofin, syrosingopine, the drug combination, and DMSO as a solvent control for 24 and 48 hours. After incubation, the cells were fixed with 50% TCA (50 µl per well) for 1 hour at 4_°_C. The plates were then washed with distilled water to remove the TCA. Subsequently, the cells were stained with 0.4% sulforhodamine B in 1% acetic acid (50 µl per well) and incubated in the dark for 30 minutes. The plates were washed with 1% acetic acid after incubation to eliminate excess SRB. After drying, the bound dye was dissolved in 10 mM unbuffered TRIS solution (150 µl per well), and the plates were shaken at 1000 rpm for 2.5 minutes. Absorbance measurements were conducted at 540 nm using an Asys UVM340 plate reader (Biochrom, Cambridge, UK). Cell viability was expressed as the percentage of living cells relative to the control (the untreated cells), calculated using the following formula:

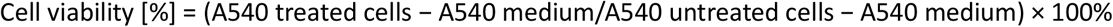

### Measurement of LDH Activity

The LDH (lactate dehydrogenase) activity was assessed using the LDH-Glo™ Cytotoxicity Assay. AsPC-1 and BxPC-3 cells were seeded and treated as previously described. Cells were treated with auranofin, syrosingopine, and their combination in a dose-dependent manner for 48 hours. After treatment, the manufacturer’s instructions were followed to evaluate the LDH activity. After incubation with LDH Detection Reagent, the luminescence signal was recorded with a GloMax® Discover reader at λ_Ex/Em_ = 495/530 nm.

### Determination of Lactate in the Medium

The lactate level in the medium was determined using the Lactate-Glo_™_ Assay following the manufacturer’s protocol. AsPC-1 and BxPC-3 cells were seeded into 12-well plates at 30 × 10^4^ cells per well and incubated for 24 hours. Subsequently, cells were treated with the tested drugs for 48h under standard conditions. After treatment, the medium was collected and incubated with Lactate Detection Reagent according to the manufacturer’s instructions. Luminescence was measured using a Lumi luminometer (Novazym, Poznan, Poland) at λ_Ex/Em_ = 495/530 nm.

### Measurement of Intracellular ATP Level in 2D Culture

The ATP level in cells was determined using the CellTiter-Glo 2.0 Cell Viability Assay following the manufacturer’s instructions. In brief, AsPC-1 and BxPC-3 cells were seeded into white 96-well culture plates (10^4^ cells/well) and, after 24h of incubation, treated with the tested compounds for 48h. Luminescence was measured using a GloMax® Discover microplate reader (Promega GmbH, Walldorf, Germany) at λ_Ex/Em_ = 495/530 nm. The intracellular ATP level was expressed as a percentage of the control cells (100%).

### Analysis of Hexokinase II Expression

To analyze HK2 expression, AsPC-1 and BxPC-3 cells were seeded into 12-well plates (30 × 10^4^/per well) and incubated for 24 h. Next, the analyzed compounds were added to the cells at appropriate concentrations and in solvent control, and the cells were incubated for 48 h. After treatment, cells were fixed for 15 minutes in 4% paraformaldehyde. The cells were then washed with PBS buffer and permeabilized for 15 minutes with 0.3% Triton-X 20 solution. Then, nonspecific binding was blocked with a 0.1 M triethylamine (TEA) solution, followed by 0.25% acetic anhydride for 5 minutes. After washing with 0.5x and 1x SSC buffer, prehybridization was performed for 1 hour in a 56°C humid chamber using the HK2 probe. Hybridization was then performed for 24 hours under the same temperature conditions, and the HK2 probe solution contained, in addition to Deinhart’s solution, EDTA, yeast tRNA, and herring single-stranded DNA (ssDNA). After completing hybridization, preparations were washed with 2x SSC, and cell nuclei were stained with DAPI dye. HK2 expression analysis was performed using an Olympus FluoView confocal microscope with FV10-ASW software (Olympus, Tokyo, Japan), and the average fluorescence in the samples was evaluated using ImageJ software.

### Evaluation of Oxidative Stress

ROS and GSH levels were measured using the ROS-Glo H2O2 Assay and the GSH-Glo Assay to assess the oxidative stress in cancer cells. AsPC-1 and BxPC-3 cells were seeded into white 96-well culture plates (10^4^ cells/well). After 24 hours of incubation, auranofin, syrosingopine, and the combination were added to the cell culture at the desired concentrations. For the ROS Assay, 6 hours before the end of incubation, the H2O2 substrate was added. After 48 hours of treatment, cells were incubated with the detection reagents (ROS-Glo and GSH-Glo, respectively). The luminescence was recorded using a GloMax® Discover microplate reader at λ_Ex/Em_ = 495/530 nm.

### Real-Time Evaluation of Cell Apoptosis

The progression of cell apoptosis was monitored using the Real-Time Glo Annexin V Apoptosis Assay. In brief, AsPC-1 and BxPC-3 cells were seeded into white 96-well plates (10^4^ cells/well) and incubated for 24h. After incubation, the appropriate concentrations of the tested compounds and a freshly prepared detection reagent were added to the cell culture, and the RealTime-Glo_™_ assay was immediately initiated. Luminescence and fluorescence were measured for 48h at specified time intervals (0, 2, 4, 6, 8, 10, 12, 24, 32, and 48h) using a GloMax® Discover at λ_Ex/Em_ = 495/530 nm.

### Assessment of Mitochondrial Membrane Potential

Mitochondrial membrane potential (MMP) was evaluated by measuring the fluorescence of the lipophilic cationic probe JC-1. AsPC-1 and BxPC-3 cells were seeded into 12-well plates (30 × 10^4^ cells/well) and incubated for 24 h. After that, the cells were treated for 48h with appropriate concentrations of the drugs. The cells were then stained with the JC-1 probe and incubated for 30 min at 37_°_C. The results were analyzed using an Olympus FluoView confocal microscope with FV10-ASW software and using FITC and TRITC filters at λ_Ex/Em_ = 514/529 nm for JC-1 monomers and λ_Ex/Em_ = 585/590 nm for aggregates. The average fluorescence in the samples was evaluated using ImageJ software.

### 3D Cell Culture (Spheroids)

Spheroids were obtained using a method based on culturing cells in a non-adhesive medium dish (liquid overlay). The procedure for spheroid preparation used 96-well culture plates coated with 1.5% agarose in RPMI medium. BxPC-3 cells were seeded into the prepared plates (15 × 10^3^ cells/well) and then centrifuged for 20 min at 10_°_C and 3000 using Sigma 3-16P (Polygen, Wroclaw, Poland). The plates were then incubated with shaking at 135 rpm for 24 h at 37°C in an atmosphere saturated with 5% CO_2_. Afterwards, 50 µL of fresh medium was added, and the cells were incubated for an additional 72 hours under normal conditions (37_°_C, 5% CO_2_). After this time, the spheroids were ready for further testing.

### Measurement of Intracellular ATP Levels in 3D Culture

To evaluate the viability of spheroids, intracellular ATP levels in the cultures were examined using the CellTiter-Glo 3D Cell Viability Assay, following the manufacturer’s protocol. The cultured spheroids were transferred to fresh white-well 96-well plates for 3D culture. Subsequently, spheroids were treated with appropriate doses of the tested drugs for 72h. Untreated spheroids served as a control. After treatment, CellTiter-Glo® 3D reagent was added to the medium to induce lysis of the spheroids. Luminescence signal was measured using a GloMax® Discover reader at λ_Ex/Em_ = 495/530 nm.

### *In Vivo* Study

The anticancer activity of auranofin and syrosingopine combination was evaluated *in vivo* using a mouse model of pancreatic cancer. Mice derived from the C57BL/6NCrL strain were intradermally implanted with 100µl of KPCY cells (200 × 10^3^ cells). After 14 days, the experimental groups were administered successively: auranofin (5 mg/kg bw), syrosingopine (5 mg/kg bw), a combination of both drugs at the same dose (5 mg/kg bw), and a solvent control (30% DMSO + 30% PEG400). Untreated mice constituted the control group. Preparations were injected into the tumor on 14, 15, 16, 17, and 18 days of the experiment. The tumor volume and body weight of the mice were monitored throughout the study. The study was terminated on the 32nd day of the experiment.

All experimental procedures were conducted following EU Directive 2010/63/EU and approved by the Bioethics Committee of the M. Sklodowska-Curie National Institute of Oncology in Gliwice. Resolution numbers: 25A/2024 (additional animals) and 25B/2024 (new or modified single activities).

### Statistical Analysis

Results are presented as mean and standard deviation (SD) values from at least three independent replicate experiments. Statistical significance analysis of differences in the obtained data and IC50 values was determined using GraphPad Prism software (Prism 10, GraphPad Software, San Diego, CA, USA). Statistical significance was calculated using a one-way or two-way ANOVA test (a modification of Tukey). Combination index (CI) values were calculated using CompuSyn 1.0 software (ComboSyn Inc., Paramus, NJ, USA).

