## Supplemental Figures 1 and 2 for "Metabolic Collapse in Pancreatic Cancer via Combined Inhibition of Lactate Export and Thioredoxin Reductase"

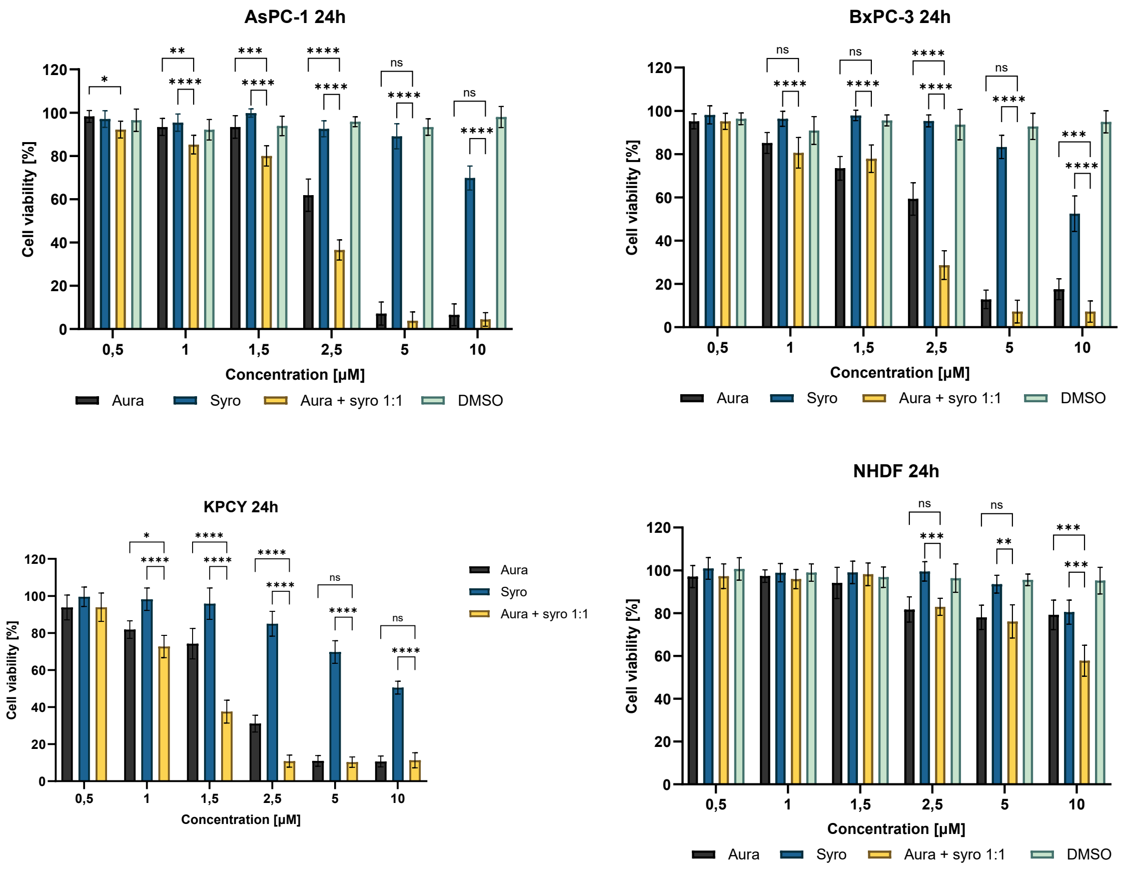


***Figure S1.*** *The viability of cells of the AsPC-1, BxPC-3, KPCY, and NHDF lines after 24-hour incubation with auranofin, syrosingopine, and a combination of both drugs at a molar ratio of 1:1. Untreated cells were the control, normalized as 100%. The number of viable cells after treatment with the test compounds was expressed as % of the control. Bar graphs show mean values with standard deviation from at least three independent biological replicates. Statistical significance was calculated using a two-way ANOVA test (Tukey's modification). ****P < 0.0001; ***P < 0.0002; **P < 0.0021; *P < 0.033; ns - not statistically significant (0.1234). Explanations of abbreviations used: Aura - auranofin, Syro – syrosingopine*

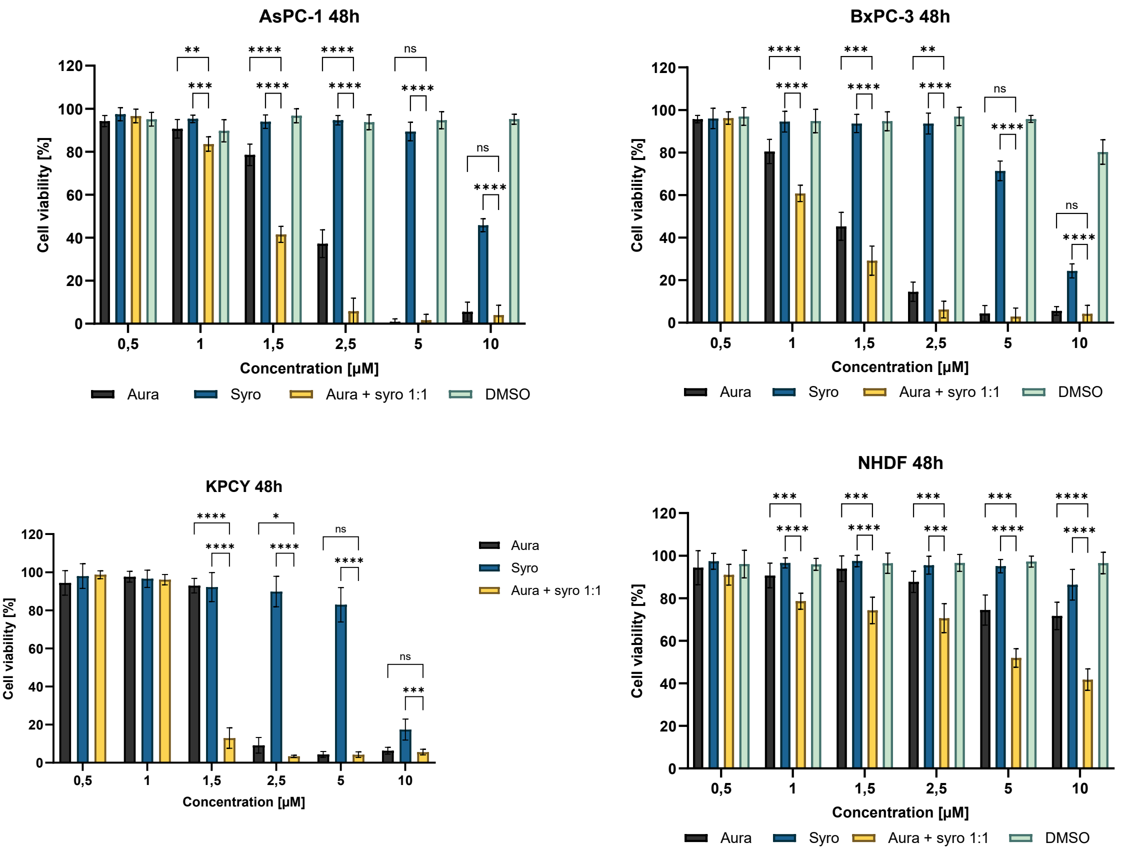


***Figure S2.*** *The viability of cells of the AsPC-1, BxPC-3, KPCY, and NHDF lines after 48-hour incubation with auranofin, syrosingopine, and a combination of both drugs at a molar ratio of 1:1. Untreated cells were the control, normalized as 100%. The number of viable cells after treatment with the test compounds was expressed as % of the control. Bar graphs show mean values with standard deviation from at least three independent biological replicates. Statistical significance was calculated using a two-way ANOVA test (Tukey's modification). ****P < 0.0001; ***P < 0.0002; **P < 0.0021; *P < 0.033; ns - not statistically significant (0.1234). Explanations of abbreviations used: Aura - auranofin, Syro – syrosingopine*
